# keju: powerful and accurate inference in Massively Parallel Reporter Assays

**DOI:** 10.64898/2026.02.24.707790

**Authors:** Albert Xue, Adam M. Zahm, Justin G. English, Sriram Sankararaman, Harold Pimentel

**Affiliations:** Bioinformatics Interdepartmental Program, UCLA, Los Angeles, CA, USA; Department of Biochemistry, University of Utah School of Medicine, Salt Lake City, UT, USA; Department of Computer Science, UCLA, Los Angeles, CA, USA; Department of Computational Medicine, David Ge!en School of Medicine, UCLA, Los Angeles, CA, USA; Department of Human Genetics, David Ge!en School of Medicine, UCLA, Los Angeles, CA, USA

## Abstract

Massively Parallel Reporter Assays (MPRAs) interrogate the regulatory function of thousands of designed genetic elements in parallel through linked DNA and RNA readouts using an engineered construct and attached minimal reporter. Given the complexity of MPRA experimental designs, several different sources of uncertainty complicate inference. We show that previous methods do not account for substantial differences in uncertainty levels between the DNA and RNA counts and between batches. Accordingly, we present `keju`, a hierarchical statistical model that estimates candidate transcription rate, differential activity between conditions, and effects from promoter composition for MPRA data. To maximize statistical power and improve false positive rate control, `keju` conditions on the DNA counts to model batch-specific and modality-specific uncertainty in the RNA counts. `keju` shows vastly improved sensitivity (59%) in simulations compared to previous methods (31% for MPRAnalyze and 9% for BCalm), and also has lower, more robust false positive rates, calling only 6.8% of unlabeled negative controls significant in real data (compared to 34% for MPRAnalyze and 12% for BCalm).

## Background

A major goal of genetics is understanding how genetic variation affects phenotypes, but heritability and GWAS hits for most complex phenotypes tend to be enriched in noncoding regions of the genome.^1–7^ Furthermore, noncoding variation is expected to influence human biology through weak effects on gene regulatory activity and the modulation of transcription, which diffuse through the system to impact higher-order processes like signaling or differentiation.^2, 8–13^ The ability to measure and perturb regulatory effects from noncoding variation in living systems, at scale, across the distribution of effect sizes, is therefore foundational to modern genetics, synthetic biology, and cell biology.

Massively Parallel Reporter Assays (MPRAs) meet this need by providing a high-throughput experimental connection between sequence and transcription (Figure 1).^14–16^ By coupling large libraries of designed genetic elements to sequenceable RNA barcodes, MPRAs link DNA and RNA counts to transform complex biological responses into high-dimensional datasets with millions to billions of perturbation-resolved observations. Once the library is introduced into cells, each construct is allowed to undergo transcription. In simpler paired designs, one plasmid pool maps to one single RNA batch.^11, 12, 17, 18^ In pooled designs, a single input plasmid pool maps to multiple RNA batches, and sometimes multiple treatments.^19–23^ In either case, these libraries can include thousands of candidates, with tens or hundreds of barcodes per candidate.^19^ After extraction, DNA and RNA sequencing are linked by barcode, and the estimand RNA /DNA is commonly used to capture regulatory activity of the corresponding candidate. Depending on design, MPRAs can also interrogate differential activity between alleles,^11, 12, 18^ between candidate responses to drug treatments,^19^ or any other comparisons.

**Figure 1.**
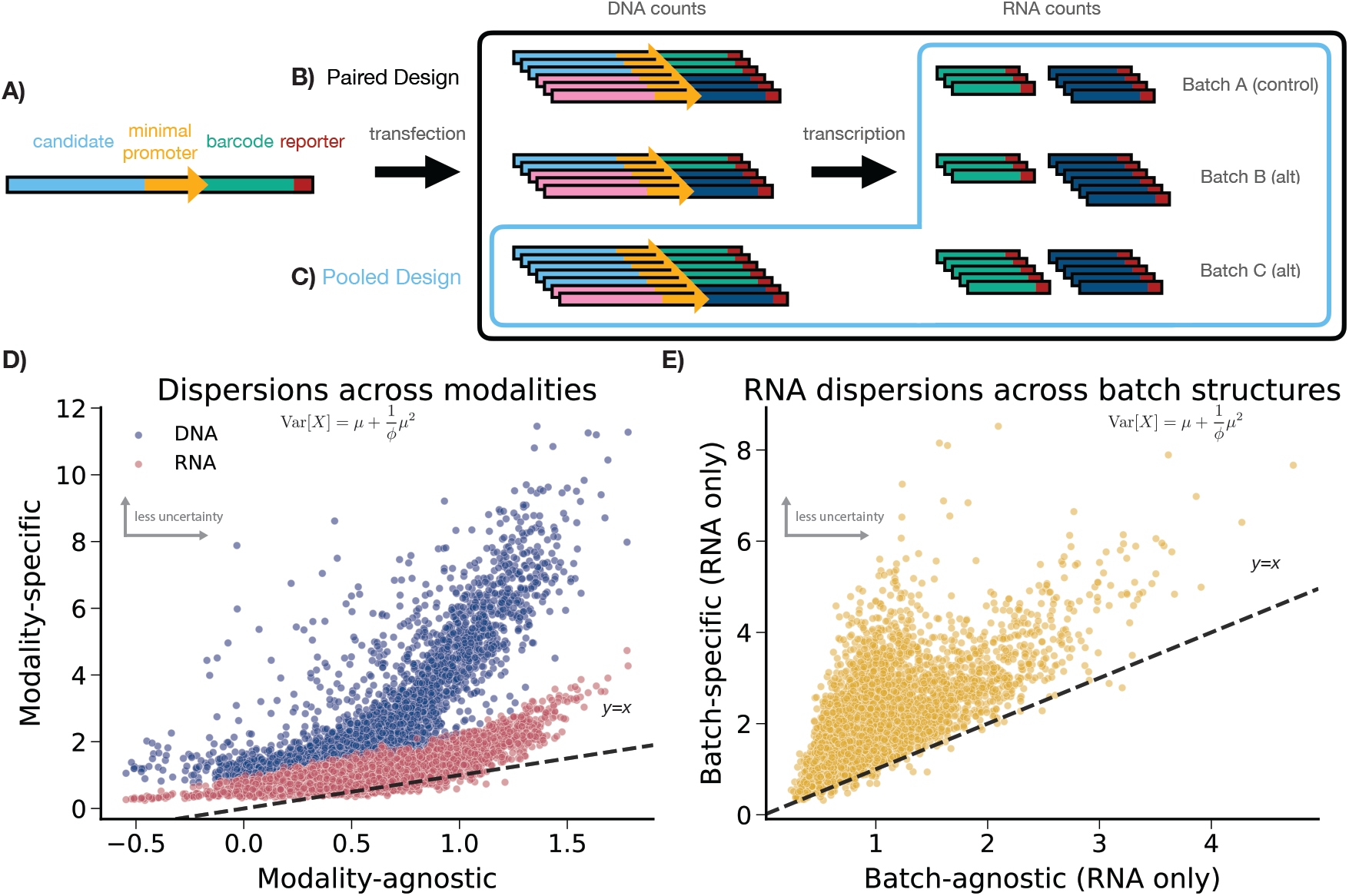
A) Standard MPRA candidates include a candidate genetic element, a minimal promoter, a unique barcode, and a reporter gene. Candidates are synthesized into a plasmid construct that is transfected into cells, which then transcribe the associated barcodes. B) In paired designs, one DNA batch maps to one RNA batch. C) In pooled designs, one DNA batch maps to multiple RNA batches. D) Each dot is an enhancer-level overdispersion estimate. X-axis is MPRAnalyze overdispersion estimate, which is modality agnostic; y-axis is modality-specific overdispersion estimates from an alternate model (detailed in Methods). Uncertainty is fundamentally larger (overdispersion estimates are smaller) in RNA counts than in DNA counts. In addition, the modality-specific model has lower uncertainty, motivating its use. E) X-axis is alternate model overdispersion estimates, y-axis is keju overdispersion estimates for a single batch. Modeling batch structure substantially improves overdispersion estimates. All data shown is PAR1_Throm.

Given the variability and complexity of MPRA experimental designs, several different sources of uncertainty complicate inference. Ashuach *et al.* previously showed that incorporating barcode-level variability substantially improves inference in MPRAs.^24^ Where previous methods like mpralm or QuASAR-MPRA summarized barcode-level information to estimate the ratio 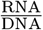, their method, MPRAnalyze, improves power by directly using barcodes as natural sources of replication.^24–26^ In addition, MPRAnalyze uses two nested Negative Binomial Generalized Linear Models (GLMs), one each on the DNA and RNA counts, to model overdispersion and the mean-variance relationship in count data.^27, 28^ The use of one GLM per modality also allows MPRAnalyze to naturally model pooled data, where competitors like mpralm and BCalm can not.^24^ However, MPRAnalyze shares a single overdispersion parameter across both GLMs, and therefore across DNA counts, RNA counts, and batches.

We argue that DNA counts, RNA counts, and batch structure (where batch structure could be related to pooling or treatment) are significant and disparate sources of uncertainty (Figure 1D,E). We further argue that separately modeling modality-specific and batch-specific uncertainty can lead to improved power, which would be important in the identification of weak effects.

In Figure 1D, we show that DNA and RNA counts in MPRAs show substantially different levels of uncertainty from each other, per-enhancer. Here, we are analyzing MPRA data from Zahm *et al.* measuring enhancer responses to activation of the PAR1 receptor by the Thrombin agonist (see Supplementary Table S1 for more). We observe that overdispersion estimates are consistently lower (uncertainty is higher) in RNA counts compared to DNA counts, and are lowest (uncertainty is highest) when sharing uncertainty estimation across both modalities as in MPRAnalyze (y=x line). The lower uncertainty in DNA counts compared to RNA counts matches the intuition that DNA counts are primarily a function of transfection, while RNA counts are downstream of transfection, transcription, and other noisy biological processes.

We also show that RNA counts also show substantial variability between batches, per-enhancer. Modeling batch-specific uncertainty in RNA counts leads to consistently higher overdispersions (uncertainty is lower) when compared to sharing uncertainty across batches (Figure 1E), indicating substantial batch-level variability. Given these results, we argue that sharing a single uncertainty estimate across these features, as in MPRAnalyze, is unintuitive given the substantial improvements observed when modeling modality-specific (Figure 1D) and batch-specific (Figure 1E) uncertainty.

Accordingly, we present `keju`, a Bayesian hierarchical model for MPRA inference that accounts for modality and batch-level differences in uncertainty quantification. `keju` is inspired by MPRAnalyze barcode-level estimation, but more closely models key design details in MPRAs. Specifically, `keju` estimates a single Negative Binomial GLM on the RNA counts only by treating DNA counts as fixed effects. As a result, `keju` models uncertainty in the RNA counts only, in a modality-specific manner. Here, we are arguing that uncertainty in DNA counts is sufficiently low to safely ignore (Figure 1D), and that uncertainty is only relevant in the RNA counts. Focusing our inference in this manner provides higher sensitivity in our benchmarking, without a corresponding tradeoff in calibration. `keju` also models batch-specific dispersions in the RNA counts to allow for variability between batches (Figure 1E).

`keju` also provides a suite of modeling features to improve inference and flexibly handle various experimental designs. To account for the mean-variance trend in count data, `keju` pools candidate enhancers with similar read coverage. In our benchmarking, mean count pooling in this fashion substantially improves `keju`‘s power to detect weak effects across datasets without substantial tradeoffs in calibration. When applicable, `keju` shrinks enhancers targeting the same motif towards motif-level estimates. This feature improves both sensitivity and calibration in nearly all our benchmarks. `keju` uses experimental negative controls to set a covariate-specific baseline behavior when estimating differential activity. This feature is helpful in the Zahm data, for example, to correct for clear differences in effect size estimates due to minimal promoter choice. Finally, for experiments using multiple minimal promoters and with motif-level shrinkage, `keju` uses motif-level information to disentangle promoter-level effects from motif-level effects. Separable estimates in this fashion allow the prediction of transcription rate for unseen promoter-motif combinations, which could be particularly important for the creation of synthetic enhancers.

Previously, MPRAnalyze was shown to have higher power and better calibration than summary-statistic methods like mpralm and QuASAR-MPRA.^24–26^ We therefore focus on benchmarking `keju` against MPRanalyze, its most direct competitor method, and BCalm, a new barcode-specific successor to mpralm that skips the barcode-level aggregation step for increased power.^29^ Details on use of MPRAnalyze and BCalm are provided in Methods.

Zahm *et al.* assess the responses of transcription factor binding motifs to stimuli that include metabolites, mitogens, toxins and pharmaceutical agents, with the eventual goal of therapeutic development and identification of drug targets. These datasets are all pooled experiments (Figure 1A), with up to 7 RNA batches per experiment but only one experiment with more than one DNA batch. From the Zahm data, we benchmark `keju`, MPRAnalyze, and BCalm using 19 case-control comparisons assessing differential activity for 6144 candidate enhancers (transcription response elements or TREs in their paper).^19^ These candidates mainly serve as replicates for 325 transcription factor binding motifs across six configurations and three minimal promoter choices, but also include 306 negative control enhancers. Since BCalm does not naturally model pooled data, we make some modifications to its usage (see Methods). Full details of batch structure and treatments are provided in Supplementary Table S1.

## Results

### Modeling MPRA-specific nuances enables robust variance estimation

We present `keju`, a hierarchical Bayesian model for MPRA analysis. For each enhancer *e*, `keju` estimates a posterior distribution on the baseline transcription rate *α_e_* in the control condition, and on the effect size *β_e_* (when applicable), marking the difference in transcription rate in the alternate condition when compared to *α_e_*. `keju` is built on four major assumptions: a.) DNA counts have sufficiently low uncertainty to be treated as fixed offsets, b.) uncertainty quantification should be batch-specific, c.) uncertainty quantification improves by sharing overdispersion estimates across enhancers with similar read coverage, and d.) enhancers targeting similar motifs have similar effects. We validate each of these assumptions in real data and simulations, and show that `keju` has better power than MPRAnalyze and BCalm in simulations while maintaining more robust false positive rate control.

The core of `keju` is a Generalized Linear Model (GLM) using a Negative Binomial distribution, parameterized such that for mean *µ*, the variance can be written as 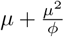 for the overdispersion parameter *ϕ*. Here, increasing *ϕ* indicates increasing certainty, and the primary advancements of `keju` come in careful treatment of the overdispersion estimates *ϕ*. The `keju` model simplifies the two-GLM Negative Binomial model in MPRAnalyze to a single GLM. Note that the Negative Binomial distribution is a common approximation for overdispersed count data.^27, 28^ With its single-GLM construction, `keju` treats DNA counts as fixed offsets, and only requires uncertainty estimation in RNA counts. Intuitively, this assumes that DNA counts have sufficiently low uncertainty that fixing them can improve power.

To justify assumption (a), Figure 1D plots modality-agnostic overdispersion estimates on the x-axis against modality-specific DNA and RNA overdispersion estimates on the y-axis. The dataset plotted is PAR1_Throm, a comparison between a control condition containing the PAR1 receptor alone, and an alternate condition containing both the PAR1 receptor and the Thrombin agonist. Modality-agnostic overdispersion estimates are from MPRAnalyze, which shares a single overdispersion for each enhancer across DNA counts and RNA counts. Modality-specific overdispersions are from an **alternate model** (not `keju`) that estimates two overdispersion per enhancer, one for DNA and one for RNA. We later use this **alternate model** as our generative model for simulations. Here, we show that DNA overdispersions are much larger (uncertainty is lower) than matching RNA overdispersions, and both are larger (uncertainty is lower) than the modality-agnostic overdispersions from MPRAnalyze. These results match our intuition that uncertainty increases as more biological processes are introduced, and emphasize that *ϕ* should be estimated per-modality.

Second, `keju` models batch structure in order to more appropriately model the variance structure, per assumption (b). Batch structure varies tremendously across MPRAs and is also frequently tied to experimental treatments and conditions. Figure 1E plots batch-agnostic overdispersion estimates against batch-specific overdispersion estimates. Batch-agnostic estimates are the modality-specific RNA estimates (y-axis) from Figure 1D, while batch-specific overdispersions are estimates from `keju` for a randomly chosen batch in the dataset. We find that batch-specific overdispersions are significantly larger than equivalent batch-agnostic overdispersions, and therefore suggest both that overdispersions should be batch-specific and that sharing overdispersions across batches likely lowers power.

Third, to incorporate assumption (c), `keju` shares overdispersion estimates across enhancers with similar counts in order to improve stability of overdispersion estimation, as in Rosace.^30^ Per batch, enhancers are grouped by average RNA count into bins of default size *G* = 50. A single overdispersion estimate *ϕ_g_* is then fit across the bin, where *g* maps (batch, enhancer) to the appropriate bin. This binning procedure helps to further account for the mean-variance trend in RNA count data, and is similar in style to the mean-variance shrinkage used by DESeq2.^27^ To assess the impact of sharing overdispersion estimates across count-grouped enhancers, we perform an ablation study by setting *G* = 1 in our benchmarking. We find that grouping overdispersion estimates in this fashion significantly increases power, with only a small tradeoff in FPR. We also find that `keju` is robust to a range of values of *G*.

Finally, per assumption (d), `keju` includes a suite of flexible priors that fit different experimental designs. Some experimental designs have multiple candidate enhancers that target the same *motif*.^19, 31, 32^ In these cases, `keju` provides motif-level regularization on both *α_e_* and *β_e_*, through motif-specific mean and variance parameters for each. We also perform an ablation study on this feature, and find that motif-level shrinkage improves both power and FPR across datasets. We also show that `keju` can use motif-level information to infer covariate-level effects on transcription rate. Users are also given the option to run `keju` without motif-level regularization.

Lastly, the Zahm data replicates candidate sequences for three different minimal promoters: human cytomegalovirus (minCMV), thymidine kinase (minTK), and the minimal promoter of Promega’s pGL4 plasmid suite (minProm).^19^ Since the goal of these experiments is the creation of synthetic promoters, we are interested in promoter-level effects on transcription rate for the same motif. We take this opportunity to extend `keju` to model promoter-specific differences on the transcription rate by jointly fitting the slope *r_k_* and the intercept *t_k_* using the motif-level mean, where *k* denotes the promoter. This feature can be reduced to motif-level shrinkage, or removed altogether. We also provide promoter-level information as covariates for the effect sizes, which have smaller promoter-level effects. Finally, we allow `keju` to set covariate-level nulls using the negative controls.

`keju` takes as input a 2 ×*N* input list 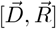 of paired barcode observations, where for each barcode *n* 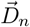 denotes the observed DNA counts and 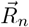 denotes the observed RNA counts. Because this processing style only requires matched barcodes, it applies equally well for both paired and unpaired data. Additional metadata denote enhancer, batch, motif, and negative controls. Only the enhancer and batch labels are strictly required -`keju` can be run without motif and control labels to fit assay design. A detailed writeup on `keju` and the alternate simulation model can be found in Methods, and a full plate model is provided in Supplementary Figure S1.

### keju has higher sensitivity in simulations

In this section, we benchmark `keju`, MPRAnalyze, and BCalm for power to detect true significant effects in simulations. To simulate data, we first fit an alternative model onto the real data, which for fairness matches neither the MPRAnalyze or `keju` models. This **alternate model** fits two Negative Binomials, one each on the DNA and RNA counts, and one overdispersion parameter is fit per-enhancer, per-modality. With two Negative Binomial GLMs, this **alternate model** is qualitatively closer to the MPRAnalyze model, but uses modality-specific uncertainty estimation (but not batch-specific uncertainty estimation). These overdispersion estimates are also used in Figure 1. For each dataset, we then draw 10 simulated data from the posterior fit of this **alternate model**, and treat the significance calls in real data fits as ground truth significance. For `keju`, we treat effect sizes with Local False Sign Rate (LFSR) less than.05 as significant, and for MPRAnalyze and BCalm we treat effect sizes with FDR less than.05 as significant.^33^ We use LFSR rather than FDR for `keju` because it more naturally summarizes the posterior distribution, and is a more stringent test than local FDR.^34^ Details are given in Methods.

In this and the following section, we use data from Zahm *et al.* In these data, a single pooled library (Figure 1B) is exposed to various treatments that range from fetal bovine serum to toxins and heavy metal compounds to GPCR activation by receptor agonists.^19^ In Figure 2 and Figure 3, the datasets are labeled in the format “{control}_{alternate} (additional information)”. Up to 100 barcodes are provided per candidate enhancer. For SF_FBS (Neuro2a cells), this comparison was performed in the Neuro-2a cell line. For GFP_ADRB2 and MRGPRX2 datasets, (A[dosage]) indicates addition of receptor agonists only, at the dosage level indicated. R indicates presence of the receptor only, and R + A[dosage] indicates presence the receptor and agonist at a specified dosage. The SF_AICAR dataset mixes multiple batches from different experimental runs, and is therefore marked with (*). GFP_ADRB2 (A[1*µ*m] <) is a subset of GFP_ADRB2 (A[1*µ*m]). Further details on datasets and abbreviations can be found in Methods and in Zahm *et al.*.

**Figure 2.**
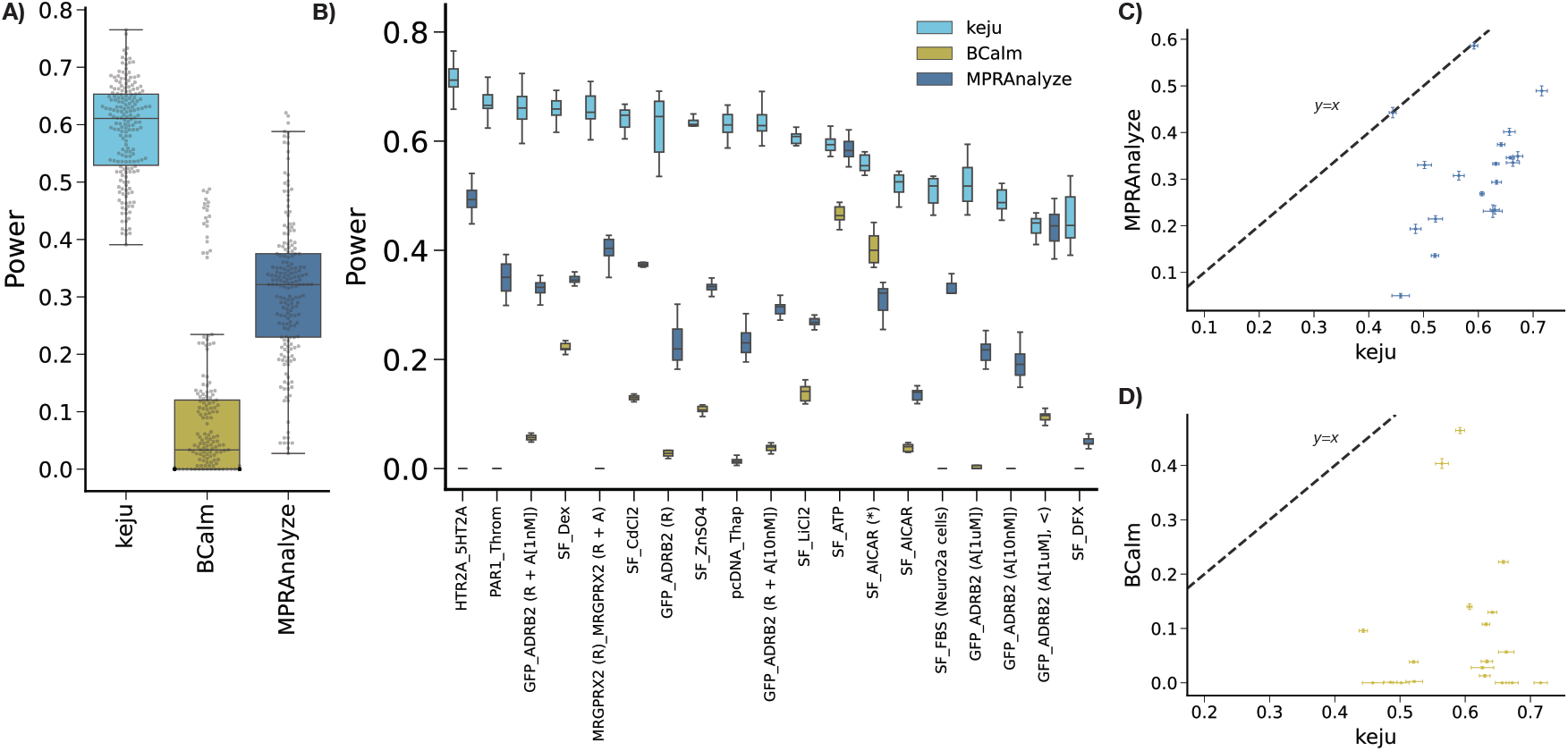
For each method, power in 10 simulations per dataset A) summarized across datasets and B) per dataset (ordered by decreasing `keju` power). Direct comparison of FPR for name against C) MPRAnalyze and D) BCalm are shown, where each point is a dataset and errorbars are standard errors.

**Figure 3.**
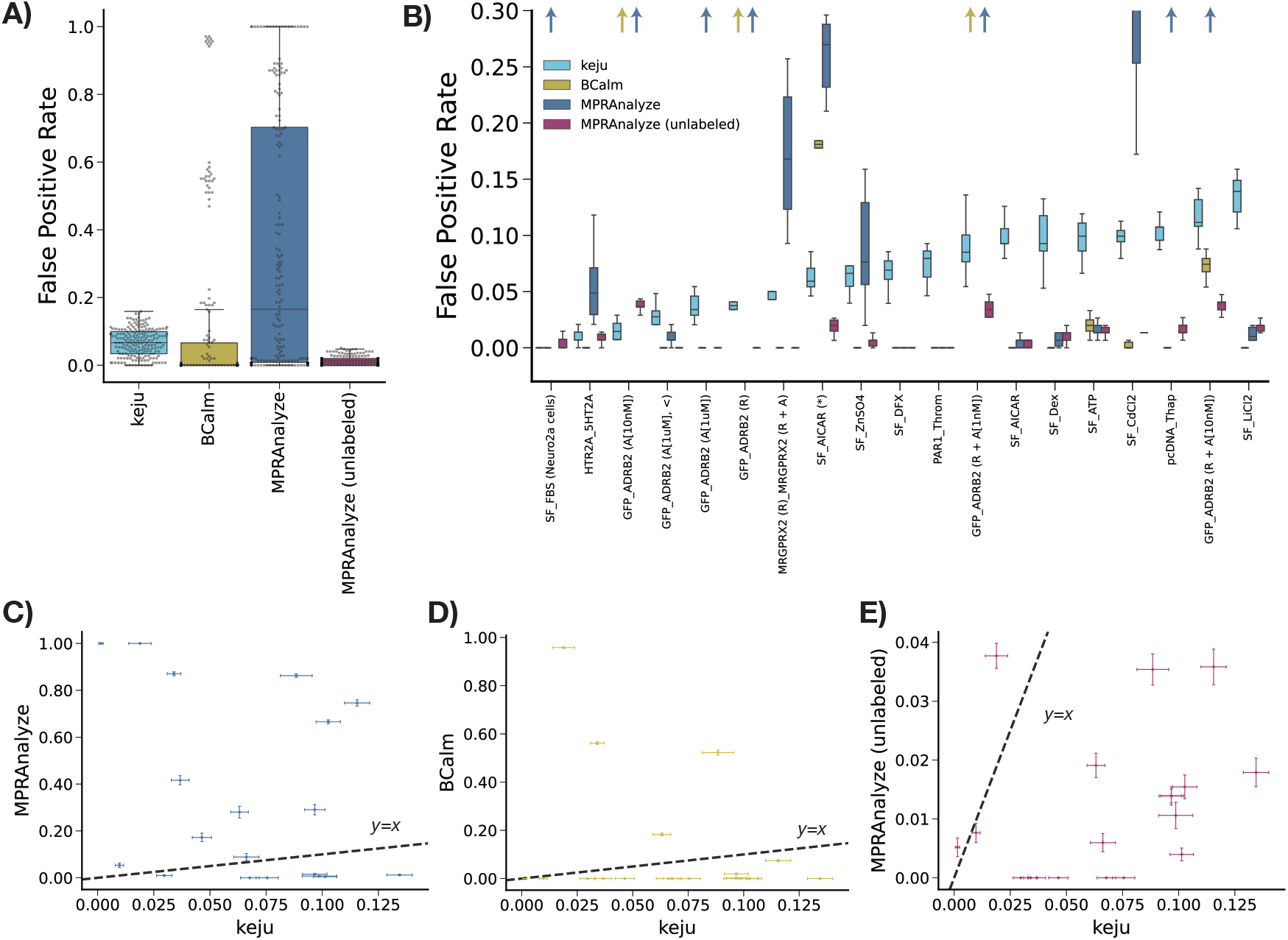
For each method, false positive rate on 10 random sets of masked negative controls per dataset A) summarized across datasets and B) per dataset (ordered by increasing `keju` FPR). Arrows represent instances where the method (indicated by color) has higher FPR than can be easily visualized. Direct comparison of FPR for `keju` against C) MPRAnalyze, D) BCalm, and E) MPRAnalyze (unlabeled) are shown, where each point is a dataset and errorbars are standard errors.

For any method, power is calculated as the proportion of ground truth significance calls that the new method calls significant. Results summarized across datasets are shown in Figure 2A, and per-dataset in Figure 2B. `keju` has substantially more power than MPRAnalyze and BCalm across all datasets (Figure 2C, D). Averaged across datasets, `keju` recalls 59.1% of ground truth significance calls compared to 31.1% for MPRAnalyze and 9.2% for BCalm.

### keju has lower, more robust false positive rate in half control experiments

In this section, we assess the false positive rate (FPR) behavior of `keju`, MPRAnalyze, and BCalm in the Zahm data. Key to this section is the use of negative controls, which are frequently scramble sequences that may have baseline transcriptional activity but should not display differential activity on average. Masking the control label serves as a proxy for assessing false positive rate (FPR). If the label of “negative control” is masked, we can treat any negative control called statistically significant as a false positive.

Both `keju` and MPRAnalyze also use these control sequences to set a control behavior, but `keju` can do so in a covariate-specific manner. For example, since the Zahm data provides three different minimal promoters per candidate enhancer, we set a separate null per minimal promoter.^19^ For BCalm, negative controls are not used to set a null, and we therefore simply calculate the FPR on the masked negative controls.

Using the Zahm data, we take the set of 306 available negative control enhancers and randomly split them in half. One half of the controls are labeled, and given to the methods as input for correction factors. The other half of the controls are masked or unlabeled, and appear to the methods as candidates. For each method, we compute the proportion of masked negative controls called significant as the FPR. We then perform this procedure across 10 random negative control maskings for each dataset.

In Figure 3, we benchmark `keju`, MPRAnalyze, “MPRAnalyze (unlabeled),” and BCalm. “MPRAnalyze (unlabeled)” is an ablation study for MPRAnalyze on controls-based correction, and is thus not provided any control labels. The FPR for each method across datasets is provided in Figure 3A, and per-dataset in Figure 3B. `keju` has lower FPR than MPRAnalyze in 12 of 19 datasets, and features substantially more stable FPR across datasets compared to both MPRAnalyze and BCalm. Averaged across draw and dataset, `keju` has 6.76% FPR, compared to 12.2% for BCalm and 34.2% for MPRAnalyze. This lower FPR is partly due to large FPR outliers for BCalm and MPRAnalyze, which in several instances call more than 20% or 50% of masked negative controls. In the real data, we also show that at comparable levels of FPR, `keju` calls more effects significant than other methods and has higher power in simulations (Supplementary Figure S2).

In Figure 3D we directly compare the FPR of `keju` and MPRAnalyze across datasets. For any dataset, we note `keju` never estimates more than 14% of masked controls significant on average, while in 6 of 19 datasets, MPRAnalyze called more than 50% the masked controls significant on average. Note that MPRAnalyze_unlabeled shows generally lower FPR than even `keju` (Figure 3E), pointing to the controls-based null as a driver of performance instability. Similarly, while `keju` has lower FPR than BCalm in only 4 of 19 datasets (Figure 3F), BCalm shows a similar instability in FPR as MPRAnalyze in three datasets. Interestingly, MPRAnalyze and BCalm share instability in the three datasets GFP_ADRB2 (A[10nM]), GFP_ADRB2 (R), and GFP_ADRB2 (R + A[1nM]). In the model fits for these data, MPRAnalyze calls 99.96%, 75.3%, and 75.9% of enhancers significant, respectively, while BCalm calls 93.9%, 31.3%, and 32.0% of enhancers significant. However, GFP_ADRB2 (A[10nM] lacks overexpression of the ADRB2 receptor, and we would therefore only expect activation of candidates downstream of endogenous ADRB2. Similarly, GFP_ADRB2 (R) overexpresses the ADRB2 receptor but does not incorporate the epinephirine agonist. As a result, we do not expect substantial global responses in either datasets.

### Motif-level shrinkage and overdispersion estimate grouping improve performance

In this section, we benchmark two ablation studies of `keju` that are viable use cases for different users. The first is “no_motif”, which removes motif-level priors for inference, and instead sets a non-informative prior for both transcription rate and effect size. The second is “no_dispersion_grouping”, which does not group overdispersion estimates by mean count (equivalent to setting *G* = 1). We find that “no_motif” and “no_dispersion_grouping” still maintain higher power and lower FPR on average than MPRAnalyze and BCalm, showing that `keju`‘s improvements on our benchmarks are robust to ablation of either modeling feature. Both ablation methods are benchmarked on the same power simulations and calibration experiments as MPRAnalyze and BCalm previously, but we separate the results into different sections for clarity. Figure 4 shows benchmarking results between `keju`, “no_motif”, and “no_dispersion_grouping”.

**Figure 4.**
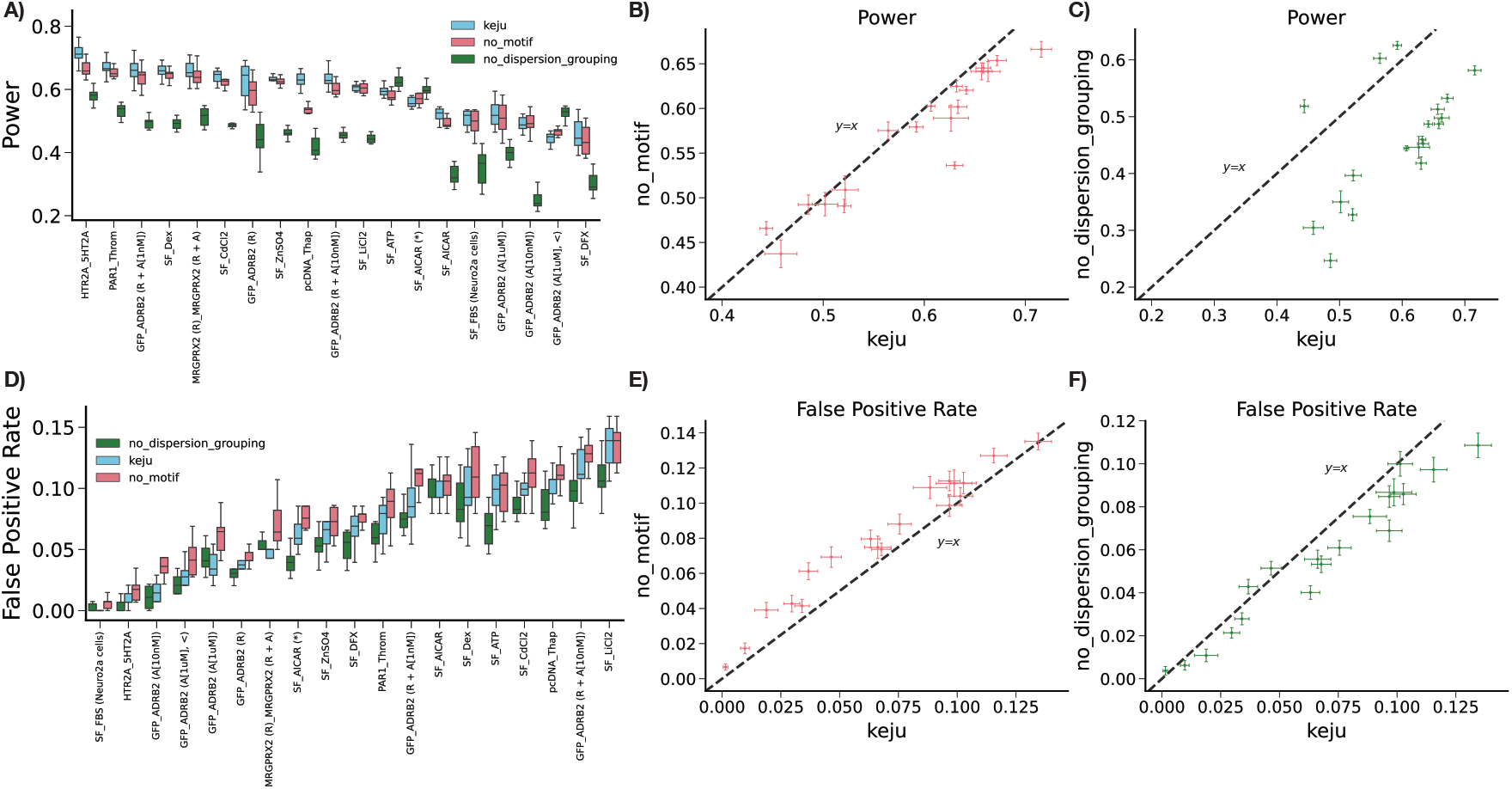
Results of ablation studies for “no_motif” (does not shrink towards motif-level mean) and “no_dispersion_grouping” (does not group overdispersion estimates). In A), we compare `keju` to both ablations in terms of power in simulations, per-dataset. In B) and C), we directly compare `keju` power in simulations to “no_motif” and “no_dispersion_grouping”, respectively. Similarly, in D) we compare `keju` to both ablations in terms of FPR. In E) and F), we directly compare `keju` FPR to “no_motif” and “no_dispersion_grouping”, respectively.

`keju` has higher power than both “no_motif” and “no_dispersion_grouping’ (Figure 4A), but also has higher FPR than “no_dispersion_grouping”. We find that the `keju` model has both higher power (Figure 4B) and lower FPR (Figure 4E) than “no_motif” on average. Performance of `keju` therefore dominates performance of “no_motif”, and we see that motif-level shrinkage in this fashion is uniformly helpful for `keju` in the Zahm data. In addition, Figure 4A and Figure 4C show that `keju` has substantially higher power than “‘no_dispersion_grouping”, suggesting that modeling the mean-variance trend in this fashion is crucial for power despite the slight FPR tradeoff (Figure 4D,F).

Supplementary Figures S3 and S4 show a full comparison of all five benchmarked methods (`keju`, “no_motif”, “no_dispersion_grouping”, MPRAnalyze, and BCalm) for power and FPR, respectively. The trends seen in Figure 2 and Figure 3 qualitatively hold when comparing ablation studies to MPRAnalyze and BCalm. For power, “no_motif” and “no_dispersion_grouping have substantially higher average power (57.2% and 45.7%, respectively) compared to MPRAnalyze (31.1%) and BCalm (9.2%). In fact, both “no_motif” and “no_dispersion_grouping” have higher power than MPRAnalyze and BCalm in all datasets except for SF_ATP, where MPRAnalyze has slightly more power than “no_motif” (Supplementary Figure S3). For FPR, “no_motif” and “no_dispersion_grouping”, like `keju`, have lower average FPR (7.9% and 5.7%, respectively) compared to MPRAnalyze (34.2%) and BCalm (12.2%), but higher average FPR than MPRAnalyze (unlabeled) (1.1%). Again, the `keju` ablation studies have lower average FPR than BCalm primarily due to the presence of a few outlier datasets where BCalm calls a high proportion of false positives (Supplementary Figure S4).

The Zahm data has a maximum of 100 barcodes per candidate enhancer, which is a high level of replication that could affect the optimal number of enhancers *G* per overdispersion estimate *ϕ_g_*. In particular, we were concerned that smaller experiments, with lower number of barcodes per candidate, would be miscalibrated at higher *G*. To understand `keju`‘s relationship to sample size and *G*, we restrict each candidate to a maximum of 10, 25, 40, 60, 80, and 100 barcodes in the real data for PAR1_Throm. We then run `keju` with *G* = 1 (“no_dispersion_grouping”), 10, 25, and 50 (base `keju`) for each subsampling level. To benchmark power, we treat significance calls in the original data as true positives and note that proportion of true positives recovered. To benchmark FPR, we recycle the negative control maskings to address FPR. Details are given in Methods.

We find `keju` has fairly robust power (Supplementary Figure S7A) and FPR (Supplementary Figure S7B) at all subsampling levels and all *G*. At most subsampling levels, the difference in power between `keju` at *G* = 10. 25. and 50 is minimal, and only separate when using all 100 barcodes per candidate (see Supplementary Figure S7A). This may be due to a substantial sample size jump when going from 80 barcodes to 100, since over 40% of candidates have more than 80 barcodes. For all *G*, power increases as *G* increases, with the biggest increase in power typically coming between *G* = 1 and *G* ≠ 1.

On the other hand, `keju` has generally consistent FPR at *G* = 10. 25. and 50 but substantially lower FPR at *G* = 1 (Supplementary Figure S7B). We find that FPR increases for all *G* until about 60 barcodes per candidate enhancer and then stabilizes. Under this threshold, `keju` FPR increases as *G* increases, but the effect is very small. Our results suggest that when grouping dispersion estimates (*G* > 1), FPR is typically stable at all sample sizes and all *G*, but for large *G* we may obtain increasing power benefits from larger sample size without trading off FPR. Finally, these results alleviate worries that `keju` would be miscalibrated in smaller experiments, even at higher *G*.

### Minimal promoter choice enables tunable transcription rate

Promoter specific modeling in `keju` allows us to examine differences between the minTK, minProm, and minCMV promoters used in the Zahm *et al.* data.^19^ We corroborate the findings that use of the minCMV minimal promoter drives higher baseline transcription rate compared to enhancers using minTK and minProm targeting the same motif (Figure 5A). For minCMV, we observe both extra baseline transcription in non-transcribed motifs, as well as a “stretching” effect for transcribed enhancers. The substantial differences in transcription rate observed for the minCMV promoter therefore render shrinkage towards a naive motif-level mean inappropriate. As a result, `keju` shrinks transcription rate estimates towards the jointly fit promoter-specific trend line 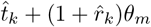, using the intercept *t_k_* and the slope 1+ *r_k_* to capture posterior distributions on these promoter-level effects on the motif-level transcription rate θ*_m_* for motif *m* (Figure 5B). Both slope estimates 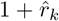 and 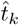 are substantially larger for the minCMV promoter, compared to minTK and minProm (Figure 5C). In contrast, the minTK and minProm minimal promoters have largely interchangeable slopes and intercepts. In general, we find these two promoters to have qualitatively similar effects in both transcription rate and significance calling. In contrast, the slope estimate from minCMV is approximately double that of minTK or minProm across datasets.

**Figure 5.**
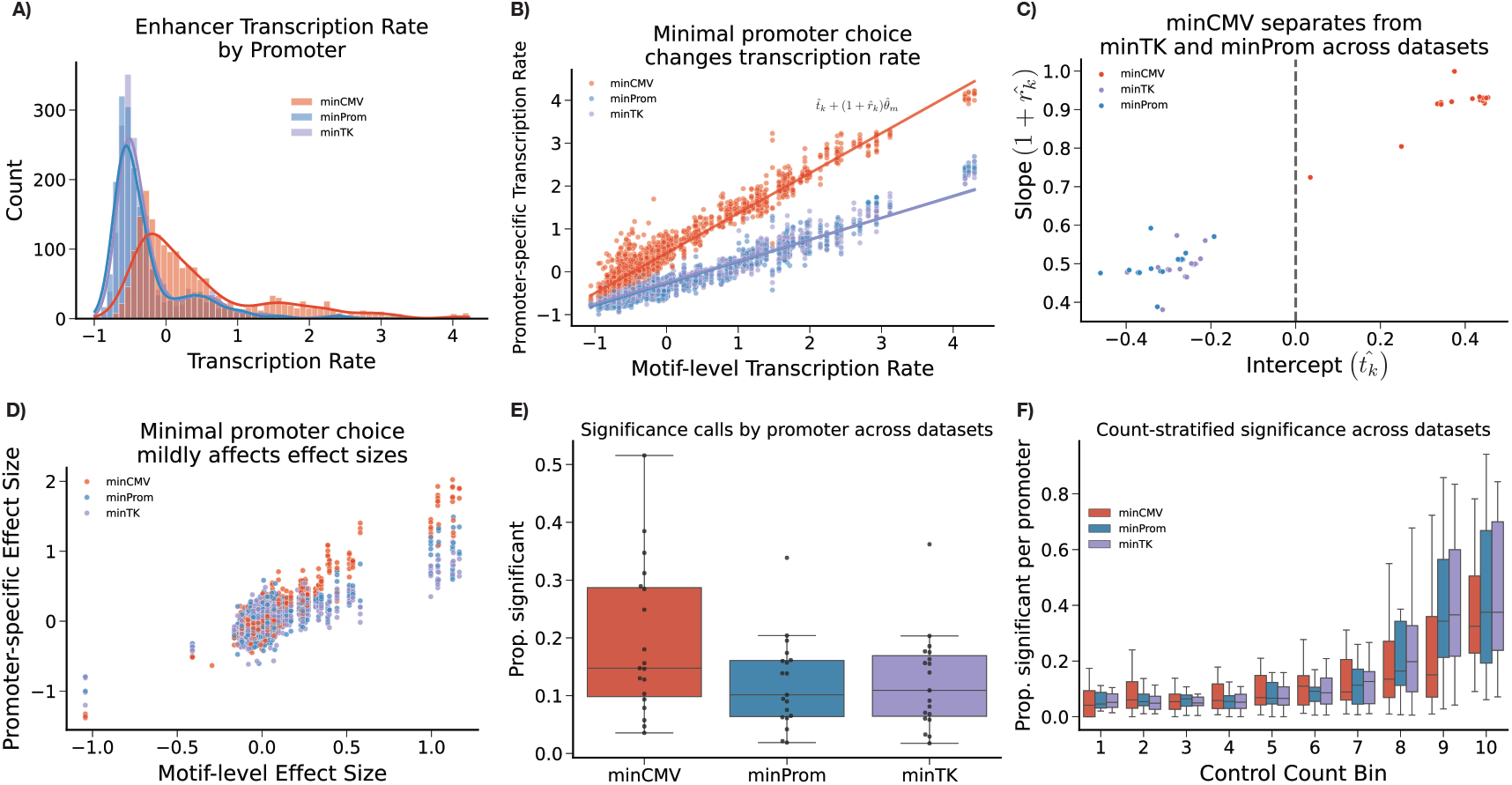
A) Transcription rate estimates vary widely depending on minimal promoter used. B) For a motif m, motif transcription rate θ*_m_* against candidate transcription rate α*_e_* for each minimal promoter. Promoter-specific intercepts 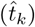 and slopes 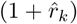 are fit by `keju`. Trend lines are 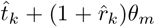. C) Intercept 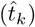 vs slope 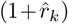 estimates for each promoter across datasets. D) Minimal promoter choice affects effect size estimates, but do not separate sufficiently to warrant separate shrinkage treatments. E) Minimal promoter choice affects significance calling for effect size estimates (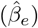). However, this effect largely disappears for `keju` F) when stratifying by counts in the control condition, across datasets. Bins are in increasing order by mean count. Boxplots are proportion of enhancers using a certain promoter called significant, across datasets. For example, the metric for minCMV enhancers in bin 1, in dataset PAR1_Throm is 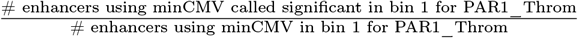. Data shown in A), B), and D) is PAR1_Throm. C), E), and F) show results across all benchmarked datasets.

A similar stretching effect can be observed in the effect sizes (Figure 5D), but is less pronounced than that of the transcription rate estimates. As a result, for the effect size estimates we do not provide promoter-specific shrinkage. Across datasets, `keju` calls a higher proportion of enhancers using minCMV significant on average (Figure 5E). To investigate this phenomenon, for each dataset we stratified enhancers into ten bins based on their mean RNA counts in the control bins. In Figure 5F, we then show the proportion of minCMV, minProm, and minTK enhancers called significant per bin, across datasets, where bin 1 has the lowest counts and bin 10 has the highest counts. For example, the metric for minCMV enhancers in bin 1, in dataset PAR1_Throm is 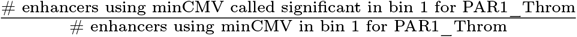 of enhancers are called significant as their average initial counts increase, regardless of promoter. However, when stratifying by bin, `keju` does not call more minCMV enhancers significant on average per bin. In fact, minCMV enhancers are called significant at a lower rate in most bins, especially in bins with higher counts. Together this suggests that, for `keju`, promoter-level effects on effect size significance calling are downstream of increased power due to increased counts, which are themselves downstream of increased transcription rate from use of minCMV.

## Discussion

We present `keju`, a Bayesian hierarchical model that improves statistical power to detect effects across the effect size distribution by prioritizing important sources of experimental variation (RNA count variance, batch structure) over less important sources (DNA count variance). `keju` thus discards uncertainty estimation on DNA counts, and estimates uncertainty only in the RNA counts. Importantly, `keju` also models variability between batches, and pools uncertainty estimation across enhancers with similar counts to model the mean-variance trend. Finally, `keju` also allows covariate correction and separate nulls per covariate to correct for known experimental biases at arbitrary granularity, and can also separate covariate-level and motif-level effects on transcription rate. In MPRAs like Zahm *et al.* where the eventual goal is the construction of sensitive enhancers whose transcription rate falls within a specified range, separate estimates of motif-level and promoter-level effects on transcription rate allow prediction of the transcription rate of unseen motif-promoter combinations. In these ways, `keju` provides an intuitive, flexible framework that improves sensitivity to detect effects, maintains robust calibration, and closely models arbitrarily complex MPRA designs.

`keju` consistently outperforms MPRAnalyze and BCalm in both power simulations and calibration experiments with masked negative controls. For calibration, `keju` improves on average FPR across datasets primarily due to its robustness. Where MPRAnalyze and BCalm frequently have datasets with large FPR outliers, `keju` shows robust FPR across all 19 datasets, and never estimates a median FPR higher than 0.15. When FPR levels are comparable, `keju` maintains higher power to detect true positive effects.

Finally, we conducted ablation studies of secondary model features like enhancer pooling and motif-level shrinkage. Both ablation studies maintain our benchmarking improvements over MPRAnalyze and BCalm, with higher sensitivity and better, more robust calibration on average across most datasets. Through these ablations, we find that motif-level shrinkage improves both sensitivity and calibration in `keju`, while enhancer pooling slightly weakens calibration but massively improves discovery power. While motif-level structure is not always present in every experiment, our results show a.) that statistical methods should take advantage of this structure when possible, and b.) that `keju` can provide powerful, reliable estimates even without this extra information.

We do note that `keju` inference can be slow since it relies on MCMC sampling, and takes around one day to run on average. Pooling enhancer overdispersion estimates reduces the number of estimated parameters, reducing runtime in extremely large datasets (Supplementary Figure S5) and substantially reducing memory usage (Supplementary Figure S6). One future improvement on `keju` could implement variational inference, especially for larger datasets.

In conclusion, we hope `keju` will provide researchers with a reliable, powerful tool for analysis of MPRA data. We expect better variance estimation through modality and batch-specific overdispersion estimates will also help downstream interpretations of enhancer effects. Moreover, our benchmarking shows that `keju` has better and more robust calibration, which then substantiates our parallel claims of higher sensitivity. As a result, we expect `keju` to confidently identify viable candidates that may have previously escaped detection, without fear of additional false positives.

## Methods

### keju model

`keju` takes as input a 2 by *N* list of DNA and RNA counts 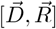, as well as additional metadata to map observations to *B_r_* RNA batches, *B_d_* DNA batches, *M* motifs, *E* enhancers, and *K* covariates. Only the enhancer and batch labels are strictly required - `keju` can be run without motif and control labels to fit assay design. Included in these *E* enhancers are *C* negative controls and *E* − *C* candidate enhancers.

For any observation *n*, essentially a tuple (*D_n_, R_n_*, barcode), these metadata map *n* to its enhancer *e*, RNA batch *b_r_*, and DNA batch *b_d_*, and motif *m*. In paired designs, the DNA and RNA batches will have a one-to-one correspondence in the metadata. In unpaired designs, the DNA batches will have a one-to-many mapping to the RNA batches, and the DNA counts will be duplicated or filled to match the RNA counts.

The binning function *g*(*e*.*b_r_*) maps the tuple (enhancer *e*, RNA batch *b_r_*) to a batch-specific mean-count bin based on the mean counts of enhancer *e* in the RNA batch *b_r_*. Per batch, all enhancers have mean RNA counts calculated and are sorted. Subsequently, the first *G* enhancers are placed in bin 1, the next *G* enhancers are placed in bin 2, etc., where by default *G* = 50. A single overdispersion estimate *ϕ_g_* is then estimated over each bin. The mapping *g* encodes the assumption that enhancers with similar counts in each batch will behave similarly, and prevents overparameterization when estimating batch-specific overdispersions.

The primary objects of interest are the enhancer-level transcription rate *α_e_* in the base condition and the enhancer-level differential activity *β_e_* for some treatment condition, compared to the base condition. For *β_e_*, we infer *K* covariates *γ_k_, k* ∈ {1.…. *K*} to correct for strong covariate-specific differences between enhancers. In the Zahm *et al.* data, we use this functionality to correct for minimal promoter choice, but this is not restricted to promoters. For identifiability, the *C* control enhancers do not estimate effect sizes, and are instead used to fit 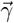. As a result, given *E* enhancers which include *C* controls and *E* −*C* candidates, we will estimate *E* transcription rates *α_e_, E* −*C* effect sizes *β_e_*, and *K* covariates *γ_k_*.

We compute RNA-batch-specific RNA normalization factors 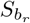 and DNA-batch-specific DNA normalization factors 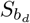. By default, we compute size factors using upper quartile normalization for both 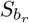 and 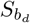, which is the same as MPRAnalyze.^24^ From 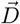 and *S_d_* we calculate the fixed offsets 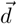, where

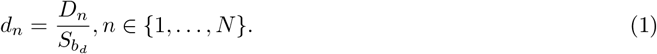

We construct the *N* × (*E* −*C*) design matrix ***X*** on the effect sizes, and the *N* ×*K* design matrix ***Y*** on the correction factors. Note that 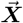 and 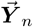 are 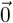 if n is an observation in the control condition. `keju` uses generalized linear regression using the Negative Binomial distribution

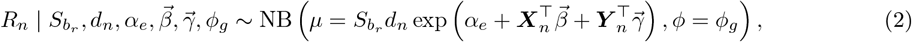

parameterized such that higher *ϕ* indicates higher certainty, i.e. 𝔼 [*R*] = *µ* and 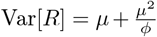. The Negative Binomial distribution is well-adapted to modeling overdispersed count data like RNA sequencing data.^27, 28^

### keju priors

The correction factors and overdispersions are given the weakly informative priors

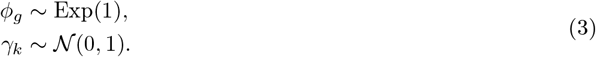

For *α_e_* and *β_e_*, we provide a suite of hierarchical shrinkage priors that flexibly model different experimental designs. We give three cases, in order of decreasing model complexity:

1. Multiple constructs target the same motif, and covariates in the experiment affect transcription.
2. Multiple constructs target the same motif, but only one promoter is used.
3. There is no motif-level structure.

For each observation *n*, case 1.) requires covariate-level and motif-level information for each observation *n*, case 2.) requires only motif-level information for each observation *n*, and case 3.) requires neither. `keju` treats *α* differently for cases 1.) and 2.). In the case of 1.), where we have a natural motif-level grouping *and* multiple promoters used in the experiment, we have the opportunity to infer promoter-specific effects on baseline construct transcription. Moreover, because of the substantial differences in observed transcription rate among enhancers using different minimal promoters (Figure 5A and Figure 5B), shrinkage towards a shared mean without including promoter-specific correction is no longer appropriate (see Figure 5C). For each motif, we infer a motif-level transcription rate contribution θ*_m_* and a motif-level variance 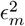. In 1.), we shrink *α_e_* to a shared motif-level mean and variance

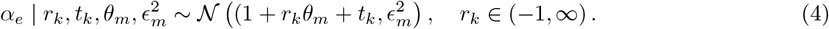

For a given motif, this model infers a promoter-specific slope *r_k_* and intercept *t_k_* across candidates targeting the same motif, and shrinks towards the motif-level mean (1 + *r_k_*) θ*_m_* + *t_k_. r_k_* is constrained to (−1. ∞) for identifiability. Both *r_k_* and *t_k_* are given the weakly informative prior

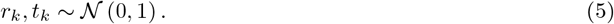

In 2.), we do not infer *r_k_* and *t_k_*. Instead, we have the simpler hierarchy on *α_e_*

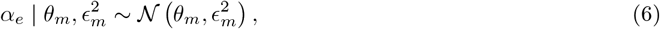

where *θ_m_* acts as the motif-level mean parameter.

In both 1.) and 2.), *θ_m_* and 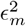 are given the weakly informative priors

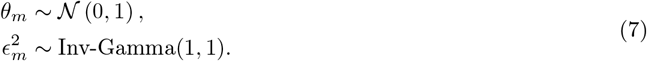

For *β*, the model is the same for cases 1.)and 2.). `keju` shrinks enhancer-level estimates of *β_e_* to a shared motif-level mean and variance

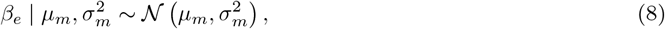

assuming that enhancers targeting the same motif will share functional effects. The mean and variance parameters are given weakly informative priors

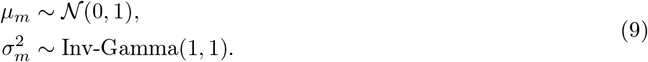

In the case of 3.), where motif-level shrinkage is not applicable, we do not infer *η_m_*.*µ_m_. ε_m_*, and *σ_m_*. Instead, *α_e_* and *β_e_* are simply given the weakly informative priors

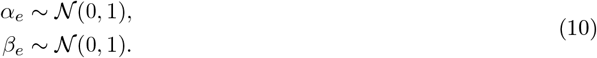

We fit `keju` with MCMC sampling in Stan using the default NUTS algorithm for Hamilton Monte Carlo.^35, 36^ Sampling was performed using 4 parallel chains, with 1000 warmup iterations and 1000 sampling iterations per chain. No divergent transitions were observed across any dataset. Sampling can take a long time to complete, dependent on the number of observations in the data.

### Data and preprocessing

We remove enhancers from data if the average number of enhancer barcodes per batch with DNA counts and RNA counts above 5 is not at least 10.

The naming convention of the datasets is “{control condition}_{alternate condition} (additional information)’. For SF_FBS (Neuro2a cells), this comparison was performed in the Neuro-2a cell line. Otherwise, this comparison was performed in the HEK293 cell line. For GFP_ADRB2 and MRGPRX2 datasets, (A[dosage]) indicates addition of receptor agonists only, at the dosage level indicated. R indicates presence of the receptor only, and R + A indicates presence the receptor and agonist. (*) indicates mixing of multiple batches from different runs. GFP_ADRB2 (A[1*µ*m] <) is a subset of GFP_ADRB2 (A[1*µ*m]). A full detailing of plasmid, treatment, dosage, and duration are provided in Zahm *et al.*^19^

The datasets benchmarked are often subsets of larger pooled experiments, and therefore frequently share common DNA batches or control RNA batches. For example, GFP_ADRB2 (A[10nm]) and GFP_ADRB2 (A[1um]) share a common DNA batch, as well as a common set of RNA batches (GFP). We provide details for batch structure in Supplementary Table S1.

### Simulations

For fairness, we use a generative simulation model separate from either `keju` or MPRAnalyze. Similarly to MPRAnalyze, this **alternate model** fits two Negative Binomial GLMs, one on the DNA counts and one on the RNA counts. However, for each enhancer *e* the simulation model fits two overdispersion estimates 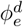and 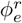, one for the DNA counts and one for the RNA counts. These overdispersion estimates in real data are also used in Figure 1. For each DNA batch *b_d_* and each enhancer *e*, we also fit a mean DNA offset 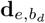. This is the only batch specific modeling that occurs, which means that this simulation model does not explicitly model batch-specific variation like `keju` does. This model also does not include motif-level shrinkage.

Because we are now performing estimation in the DNA count space, rather than treating them as fixed offsets, we must treat the input differently. While `keju` takes in a 2 by *N* list of DNA and RNA counts 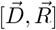, and is not sensitive to duplication of DNA batches or counts as long as barcodes match, this input format would inflate overdispersion estimates in the simulation model. We thus provide our simulation model a list of DNA counts 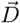 of length *n_d_*, where *n_d_* denotes the number of counts in the DNA batch. 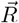 is a list of RNA counts of length *n*, as before, where now *n_d_* ≠ *n*. The model then becomes

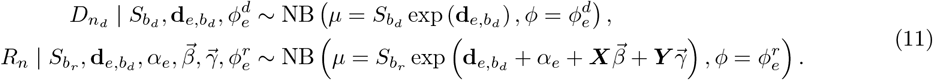

We give noninformative priors for each parameter

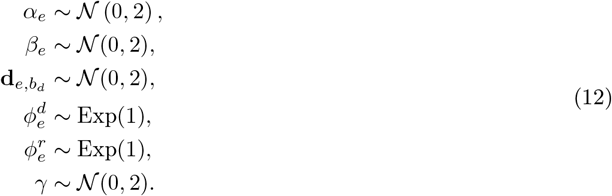

To draw simulations, we first fit the simulation model onto real data and then draw simulations from the posterior fit. We fit this model onto all 19 datasets benchmarked using MCMC in Stan,^35^ and draw ten simulations for each dataset. For each simulation, we randomly choose a draw in the posterior fit, and draw data from the same generative model (Eq. 11). Simulations mirror the batch structure of their original dataset. If an enhancer has less than 25 barcodes in the original data, we draw the same number of barcodes for that enhancer. Otherwise, we cap the maximum number of barcodes per enhancer at 25. We also add a pseudocount of 1 to the DNA counts.

### MPRAnalyze

We ran MPRAnalyze in two fashions. We run generic MPRAnalyze by providing the object control labelings. This approach is represented as “MPRAnalyze” in the manuscript. In Figure 3, “MPRAnalyze_unlabeled” is run without control labelings, but all other details are the same.

The way we run MPRAnalyze is also different for SF_AICAR (*) than any other dataset, because this dataset uses one DNA batch for control treatment RNA batches, and a separate DNA batch for alternate treatment RNA batches. This design is because SF_AICAR (*) tests the use of batches from different runs. For other datasets, we perform depth factor estimation on DNA counts without library size factors, and perform depth factor estimation on RNA counts using batch and condition as library size factors. For SF_AICAR (*), we include condition as a library size factor for the DNA counts, which is equivalent to using batch.

The regression designs for the DNA regression are also different. For *analyzeQuantification*, we use the dna design ∼ 1 for most datasets, and ∼*condition* for SF_AICAR (*). For *analyzeComparative*, we use the dna design ∼*barcode* for most datasets, and ∼*barcode* + *condition* for SF_AICAR (*). The RNA designs are universally ∼ condition, and the reducedDesign is universally ∼ 1. Significance calls are based on adjusted p-values at a threshold of 0.05. We run MPRAnalyze with 12 cores.

### BCalm

BCalm assumes that barcodes have a one-to-one mapping from DNA batch to RNA batch, and further that each pair of (DNA, RNA) batches shares a treatment. As a result, it does not naturally handle pooled data. To accommodate these expectations, we duplicate the barcodes in the experiment so that each barcode has a “REF” and “ALT” version. Similarly, for each (RNA batch, treatment condition) pair in the data, we duplicate the shared DNA counts. NaNs fill duplicated space where the experiment has no data. We then run run bcalm, or equivalently mpralm, with the parameters aggregate=“none”, normalize=TRUE, model_type = “corr_groups”, and plot=FALSE. Significance calls are based on adjusted p-values at a threshold of 0.05.

### Subsampling Experiments

In Figure S7, we re-use the data from the half controls experiments for PAR1_Throm. We do this to present replicability on false positive rate estimates, but there is no true sense of replicability for power since all the data is the same (only the maskings change). As a result, Figure S7 does not have standard errors.

In this data, and the Zahm data in general, barcodes are numbered from 1 to 100. For each replicate, we run `keju` with *G* = 1. 10. 25. and 50. To assess power, we use the significance calls of `keju` at *G* = 50 in the original data as ground truth. Then, in Figure S7A, power is the proportion of original significance calls called significant at each subsampling point, for each *G*. Note that at 100 barcodes with *G* = 50, `keju` recovers 100% of the original significance calls. To assess false positive rate, we perform the same procedure as in Figure 3, where FPR is the proportion of masked negative controls called significant. Since we have multiple random maskings, Figure S7B does have a notion of distribution spread.

## Declarations

### Ethics approval and consent to participate

Not applicable.

### Consent for publication

Not applicable.

### Availability of data and materials

`keju` is available to run as an R package and can be installed from https://github.com/pimentellab/ `keju`. Code and data for all the analyses in this paper can be found at https://github.com/asxue/ `keju`-paper-analysis.

### Authors’ contributions

AX, JGE, SS, and HP conceived the project. AX, SS, and HP developed the statistical model. AX, SS, and HP designed the simulations. AX performed benchmarking and data analysis, with input from AMZ, JGE, SS, and HP. AX, SS and HP wrote the manuscript, with input from AMZ and JGE. All authors read and approved the final manuscript.

## Acknowledgements

The authors thank the HP and SS labs for their many stimulating discussions. This work used computational and storage services associated with the Hoffman2 Cluster which is operated by the UCLA Office of Advanced Research Computing’s Research Technology Group.

