## supplement for "keju: powerful and accurate inference in Massively Parallel Reporter Assays"

### S1 Supplementary Figures

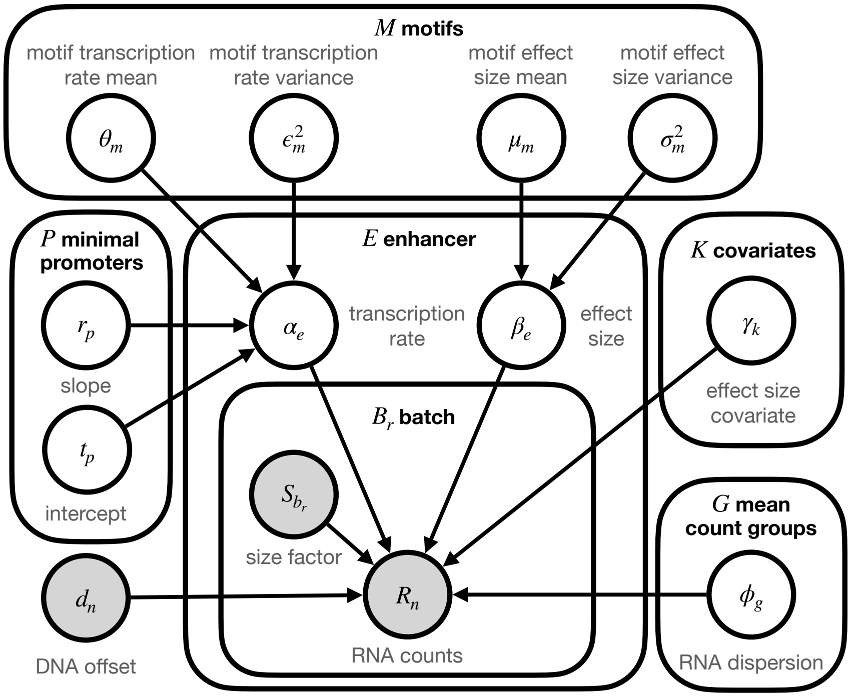


Figure S1: Bayesian plate model representation of `keju`. Distributional details are in Methods.

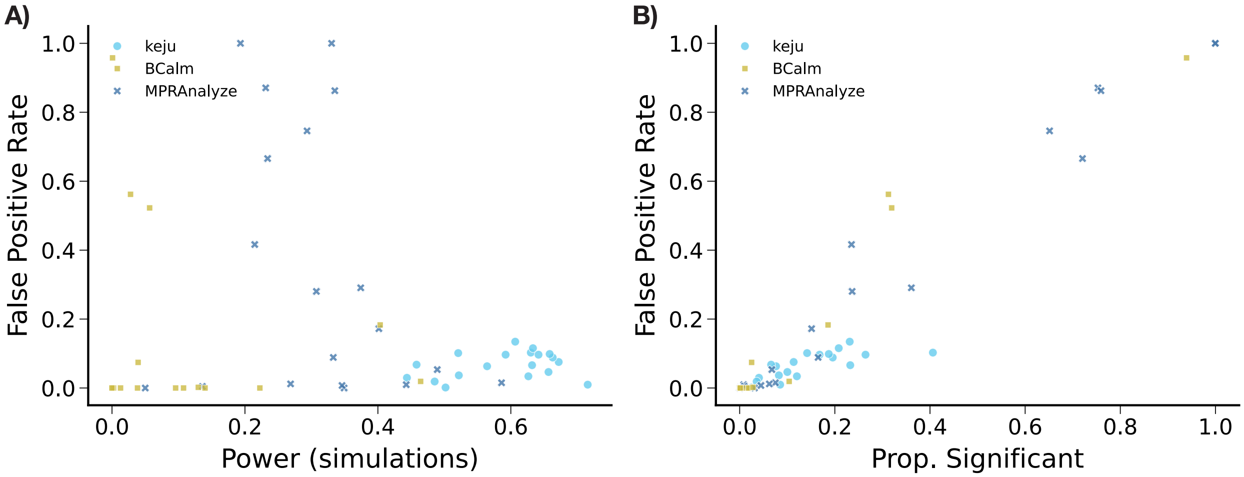


Figure S2: For all methods and datasets, A.) power in simulations and B.) proportion of effects called significant in real data, against average false positive rate in half control experiments. Each dot is a dataset.

| Dataset | DNA batches | Control RNA batches | Control Condition | Treatment RNA batches | Treatment Condition |
|---|---|---|---|---|---|
| GFP_ADRB2(A[1uM], <) | 1 | 2 | none | 1 | Epinephrine (1uM) |
| SF_AICAR (*) | 2 | 4 | none | 1 | AICAR (100 uM) |
| SF_ATP | 1 | 4 | none | 3 | ATP/GTP (50 uM) |
| HTR2A_5HT2A | 1 | 1 | HTR2A receptor | 1 | Serotonin (100nM) |
| PAR1_Throm | 1 | 1 | PAR1 receptor | 1 | Thrombin (10nM) |
| MRGPRX2 (R)_ MRGPRX2 (R + A) | 1 | 1 | MRGPRX2 receptor | 1 | (R)-zn3573 (3uM) |
| SF_FBS (Neuro2a cells) | 1 | 1 | none | 1 | Fetal bovine serum (10%) |
| pcDNA_Thap | 1 | 2 | none | 1 | Thapsigargin (2uM) |
| GFP_ADRB2 (A[10nM]) | 1 | 2 | none | 1 | Epinephrine (10nM) |
| GFP_ADRB2 (A[1uM]) | 1 | 2 | none | 1 | Epinephrine (1uM) |
| GFP_ADRB2 (R) | 1 | 2 | none | 1 | ADRB2 receptor |
| GFP_ADRB2 (R + A[10nM]) | 1 | 2 | none | 1 | ADRB2 receptor + Epinephrine (1uM) |
| GFP_ADRB2 (R + A[1nM]) | 1 | 2 | none | 1 | ADRB2 receptor + Epinephrine (1nM) |
| SF_ZnSO4 | 1 | 4 | none | 1 | ZnSO4 (200 uM) |
| SF_DFX | 1 | 4 | none | 1 | Deferoxamine (100uM) |
| SF_LiCl2 | 1 | 4 | none | 1 | LiCl2 (40mM) |
| SF_AICAR | 1 | 4 | none | 1 | AICAR (100 uM) |
| SF_Dex | 1 | 4 | none | 1 | Dexamethasone (100nM) |
| SF_CdCl2 | 1 | 4 | none | 1 | CdCl2 (30 uM) |

Table S1: Description of batch structure, control conditions, and treatment conditions for all datasets. For SF_AICAR (*), data from multiple experimental runs are used. One set of data is used for the control condition, and another set of data is used for the treatment condition. More information can be found in Zahm *et al.*, Supplementary Data 1.

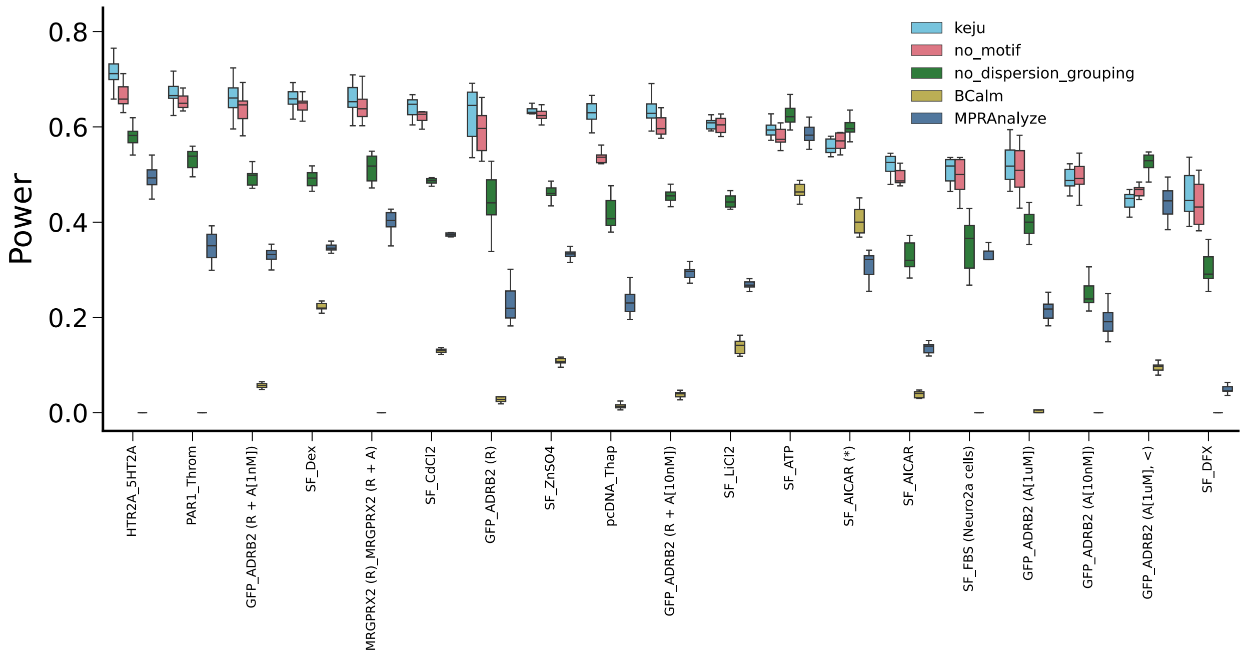


Figure S3: Power in simulations, for all methods and ablations.

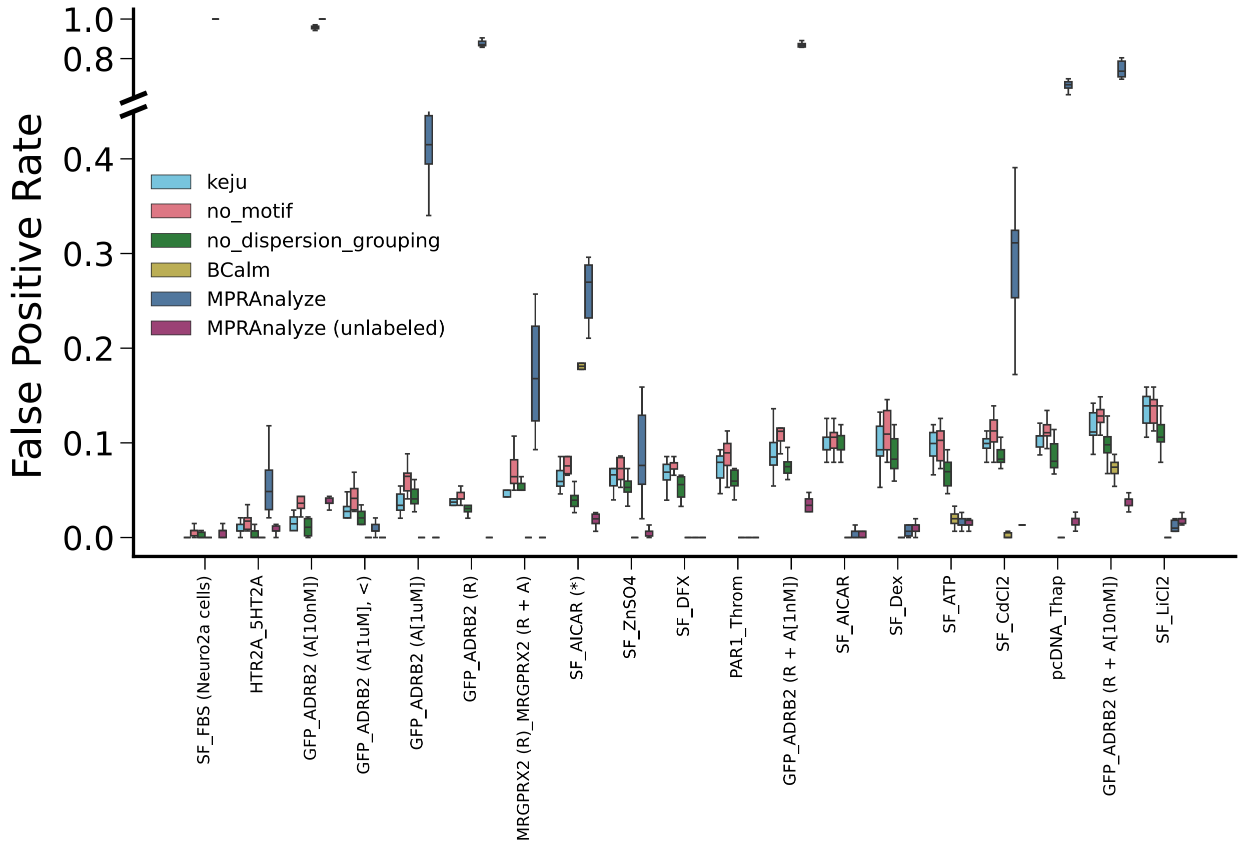


Figure S4: False positive rate in half-control experiments, for all methods and ablations. Note that y-axis is broken from 0.45 to 0.6 to accomodate MPRAnalyze and BCalm.

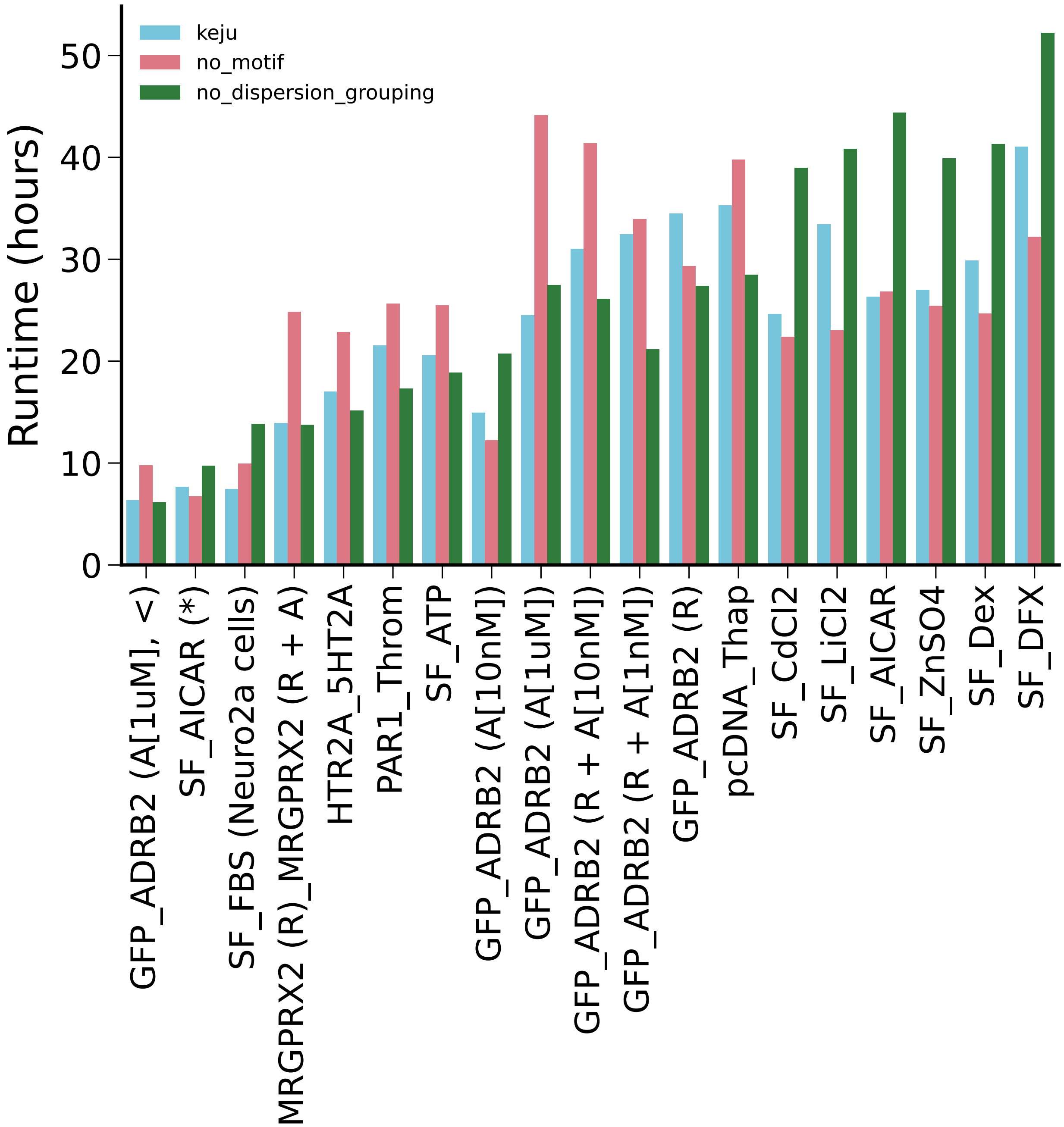


Figure S5: Runtime for each method with parallel sampling across four cores. Datasets are ordered in terms of increasing number of observations (number of barcodes multiplied by number of RNA batches), which ranges from 334776 to 1963980.

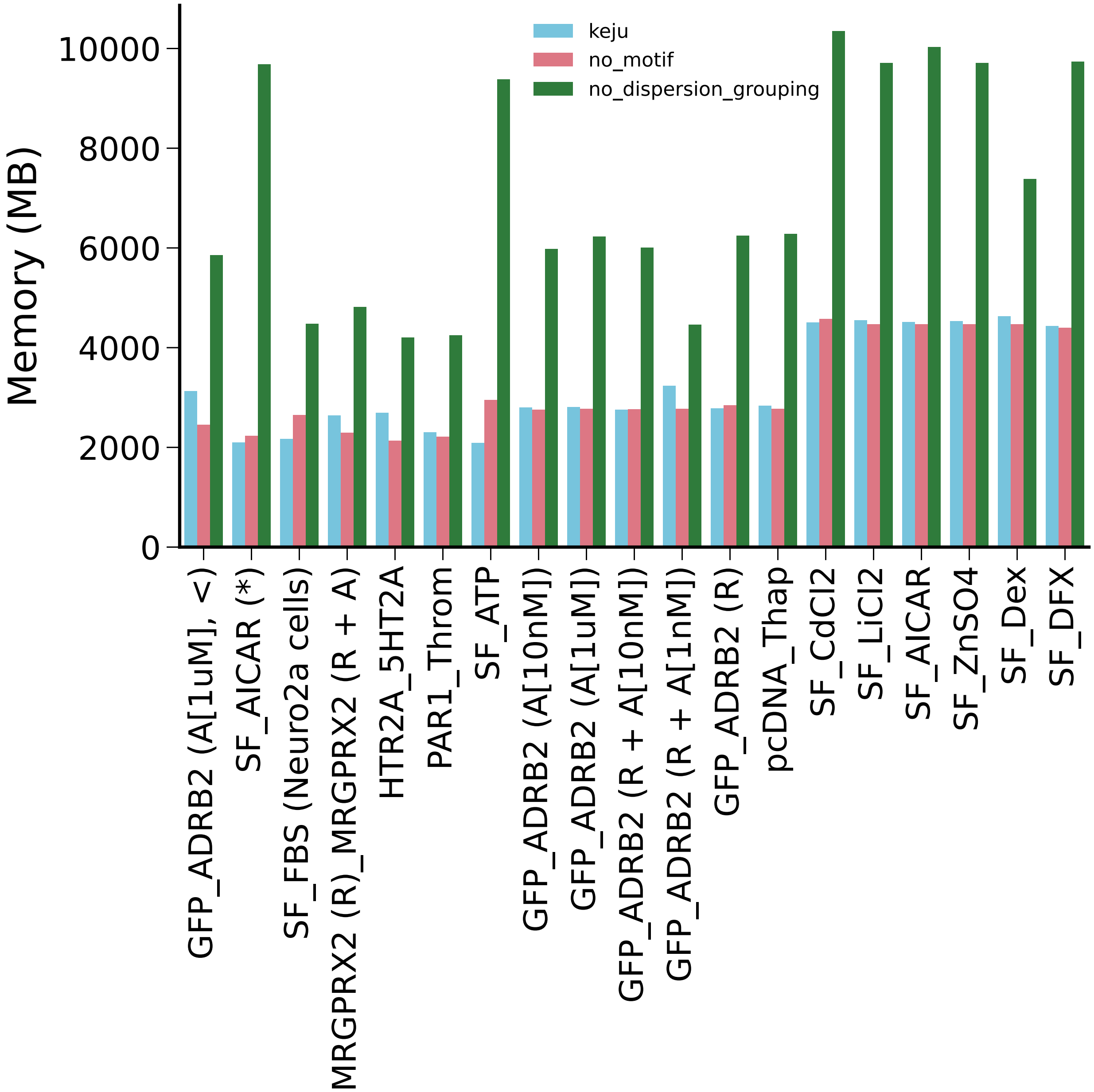


Figure S6: Memory usage of each method across four cores. Datasets are ordered in terms of increasing number of observations, which ranges from 334776 to 1963980.

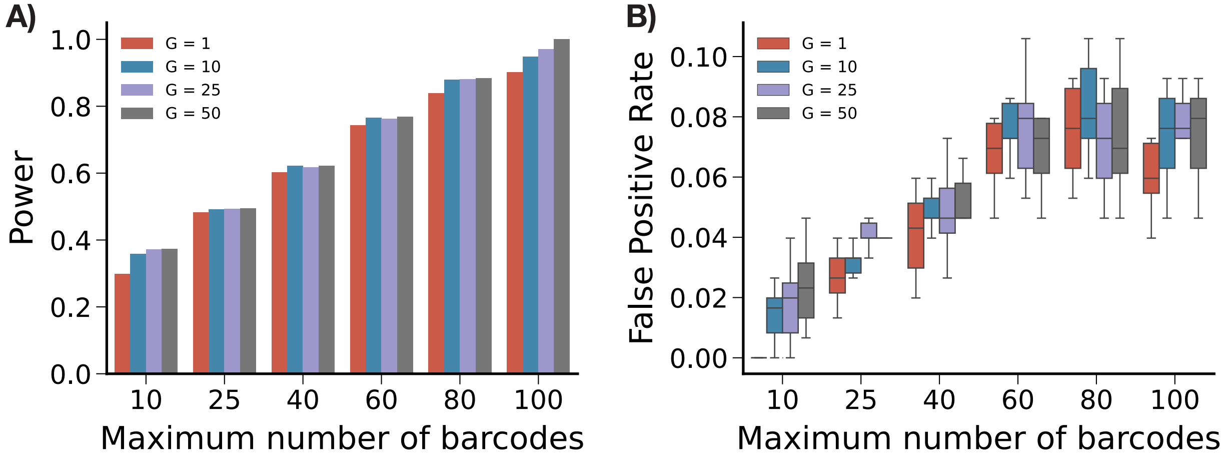


Figure S7: Maximum number of barcodes allowed per enhancer in subsampling experiment against A) power, measured as proportion of significance calls recovered from original fit with all barcodes, and B) false positive rate, measured as proportion of masked negative controls called significant. All data shown is PAR1_Throm.
